# Generating synthetic signaling networks for in silico modeling studies

**DOI:** 10.1101/2020.05.08.084848

**Authors:** Jin Xu, H Steven Wiley, Herbert M Sauro

## Abstract

Predictive models of signaling pathways have proven to be difficult to develop. Traditional approaches to developing mechanistic models rely on collecting experimental data and fitting a single model to that data. This approach works for simple systems but has proven unreliable for complex systems such as biological signaling networks. Thus, there is a need to develop new approaches to create predictive mechanistic models of complex systems. To meet this need, we developed a method for generating artificial signaling networks that were reasonably realistic and thus could be treated as ground truth models. These synthetic models could then be used to generate synthetic data for developing and testing algorithms designed to recover the underlying network topology and associated parameters. We defined the reaction degree and reaction distance to measure the topology of reaction networks, especially to consider enzymes. To determine whether our generated signaling networks displayed meaningful behavior, we compared them with signaling networks from the BioModels Database. This comparison indicated that our generated signaling networks had high topological similarities with BioModels signaling networks with respect to the reaction degree and distance distributions. In addition, our synthetic signaling networks had similar behavioral dynamics with respect to both steady states and oscillations, suggesting that our method generated synthetic signaling networks comparable with BioModels and thus could be useful for building network evaluation tools.

**Highlights:**

- We provided a Julia script to generate synthetic signaling networks.
- We defined reaction degree and distance to measure the reaction network topology.
- We provided the Python scripts to calculate the reaction network topology.
- The synthetic signaling networks had topological similarities with the BioModels.
- The synthetic signaling networks had dynamic similarities with the BioModels.

## 1. Introduction

In this article, we described a script in the modern language of Julia [1] that could be used to generate synthetic signaling networks. Such networks could be used as a basis to test novel algorithms designed to identify the topology and parameters of real signaling networks from biological data. This is particularly important for building predictive models of signaling networks involved in diseases. If reliable predictive dynamic models of signaling pathways could be developed, then more rational drug design and targeting would be possible. However, developing predictive mechanistic models of signaling pathways is difficult due to the large number of interactive components (including potentially unknown interactions) and the sparsity of suitable data to calibrate them. To address this need, we have been developing perturbation-based approaches to infer the underlying topology of signaling networks. Synthetic networks would be useful in this effort, but they must recapitulate the biophysical constraints that govern signaling pathways. Thus, we set out to define a general approach to generating synthetic signaling networks that could appropriately resemble natural ones.

Biological signaling pathways [2] are information-processing networks that are used by cells to translate external signals and cues into appropriate cell actions, such as cell growth, differentiation, movement, death, or metabolic activity. Signaling pathways are based on the interaction of a network of proteins through a limited number of processes, such as phosphorylation, protein complex formation, and targeted cleavage, degradation, or synthesis [3]. A cell’s response to external signals is typically mediated by specific receptor proteins, which carry the information into cells through a series of complex steps that amplify and process the signal before its output to the effector function. The amplification and signal processing steps frequently use enzymatic steps, such as protein phosphorylation or proteolysis. Signaling pathways also frequently display multiple feedback loops that regulate the dynamic behavior of the network and its information-processing ability. In many cases, the role of feedback loops and the overall topology of the signaling pathways is unclear.

Methods for developing predictive models of biochemical pathways still have many limitations. Many of the commonly used techniques have been translated directly from other disciplines where systems tend to be much simpler. One approach is to collect experimental data and fit it to a single model using a suitable optimization algorithm [4, 5]. Given the large number of state variables and parameters present in signaling network models, the availability of limited data results in severe overfitting and non-identifiability of parameters [6, 7], even assuming that the topology of the model is correct. Such models fail to generalize and often have poor predictive value. This problem is by no means restricted to just biological systems but applies to any complex system, such as weather forecasting [8], climate models [9, 10], financial models [11] or hydrodynamic models [12]. Given the difficulties and importance of being able to develop predictive models of such systems, there is a general need for the development of novel approaches that take into account the uncertainties in our knowledge and ability to make measurements that will generate models with the most predictive power. For example, the silico evolution algorithm in biochemical networks [13] is applicable to create several oscillators or bistable switches [14]. The differential algorithm [15] is applicable for parameter optimization search for reaction networks [16]. However, evaluating the effectiveness of any new model generating or new algorithm requires the availability of “ground truth” models against which algorithm output can be compared. One approach is to generate artificial signaling networks that are reasonably realistic and can serve as ground truth models. Such models can be used to generate artificial “experimental” data that can be used to test an algorithm’s ability to recover the original model and species parameters. Especially in the era of Artificial Intelligence (AI), the generated synthetic signaling networks could provide sufficient training data or a baseline for validation.

In the last 60 years, tools such as BioNetGen [17], Systems Biology Toolbox for Matlab [18], COPASI [19], PySB [20], Tellurium [21, 22], etc. have greatly contributed to model biological systems. In addition, there is work regarding network inference methods helping understand multi-omics [23], and there are related tools such as Causal-Path [24] and MAGICIAN [25]. In this article, we described a computational approach for generating synthetic signaling networks, which used reaction motifs [26] as a way to populate the biochemical reaction networks. In addition, we also did topological analysis for the generated synthetic signaling networks, which was a further contribution to the existing methods of biological network inference, compared with downloaded signaling networks from the BioModels Database. Our work could also enable novel interaction exploration using the other existing aforementioned tools with the popular output format of Systems Biology Markup Language (SBML) [27].

## 2. Methods

To generate synthetic networks, we first created a list of unit processes that are to be found in most signaling networks. These were shown in Figure 1, including catalyzed transformations (A), three binding and unbinding reactions (B-D), and two phosphorylation/dephosphorylation units (E, F) which included single (E) and doubly-phosphorylated (F) motifs. The three binding and unbinding reactions (B-D) followed mass-action kinetic rate laws. The other three units (A, E, F) followed reversible Michaelis-Menten rate laws. Their corresponding rate equations were shown in Table 1. All rate constants and species concentrations were assigned randomly. The single-cycle case used two reversible Michaelis-Menten kinetics for both phosphorylation and dephosphorylation. For the double circle case, we made use of four reversible Michaelis-Menten kinetics, which was a special case of the single cycle.

**Table 1.** The rate equations of the six types of units composing the artificial random networks. The three binding and dissociation reactions (B-D) followed mass-action kinetic rate laws. The other three units (A, E, F) followed reversible Michaelis-Menten rate laws.

| Unit type | Rate laws |
| --- | --- |
| uni-uni (A) | $v_A = \frac{C(k_f \cdot A/K_A - k_r \cdot B/K_B)}{1 + A/K_A + B/K_B}$ |
| uni-bi (B) | $v_B = k_f \cdot A - k_r \cdot B \cdot C$ |
| bi-uni (C) | $v_C = k_f \cdot A \cdot B - k_r \cdot C$ |
| bi-bi (D) | $v_D = k_f \cdot A \cdot B - k_r \cdot C \cdot D$ |
| Single phosphorylation<br>dephosphorylation cycle (E) | $v_{E1} = \frac{C(k_{f1} \cdot A/K_{A1} - k_{r1} \cdot B/K_{B1})}{1 + A/K_{A1} + B/K_{B1}}$ $v_{E2} = \frac{D(k_{f2} \cdot B/K_{B2} - k_{r2} \cdot A/K_{A2})}{1 + B/K_{B2} + A/K_{A2}}$ |
| Dual phosphorylation<br>dephosphorylation cycle (F) | $v_{F1} = \frac{D(k_{f1} \cdot A/K_{A1} - k_{r1} \cdot B/K_{B1})}{1 + A/K_{A1} + B/K_{B1}}$ $v_{F2} = \frac{D(k_{f2} \cdot B/K_{B2} - k_{r2} \cdot C/K_{C2})}{1 + B/K_{B2} + C/K_{C2}}$ $v_{F3} = \frac{E(k_{f3} \cdot C/K_{C3} - k_{r3} \cdot B/K_{B3})}{1 + C/K_{C3} + B/K_{B3}}$ $v_{F4} = \frac{E(k_{f4} \cdot B/K_{B4} - k_{r4} \cdot A/K_{A4})}{1 + B/K_{B4} + A/K_{A4}}$ |

**Figure 1:**
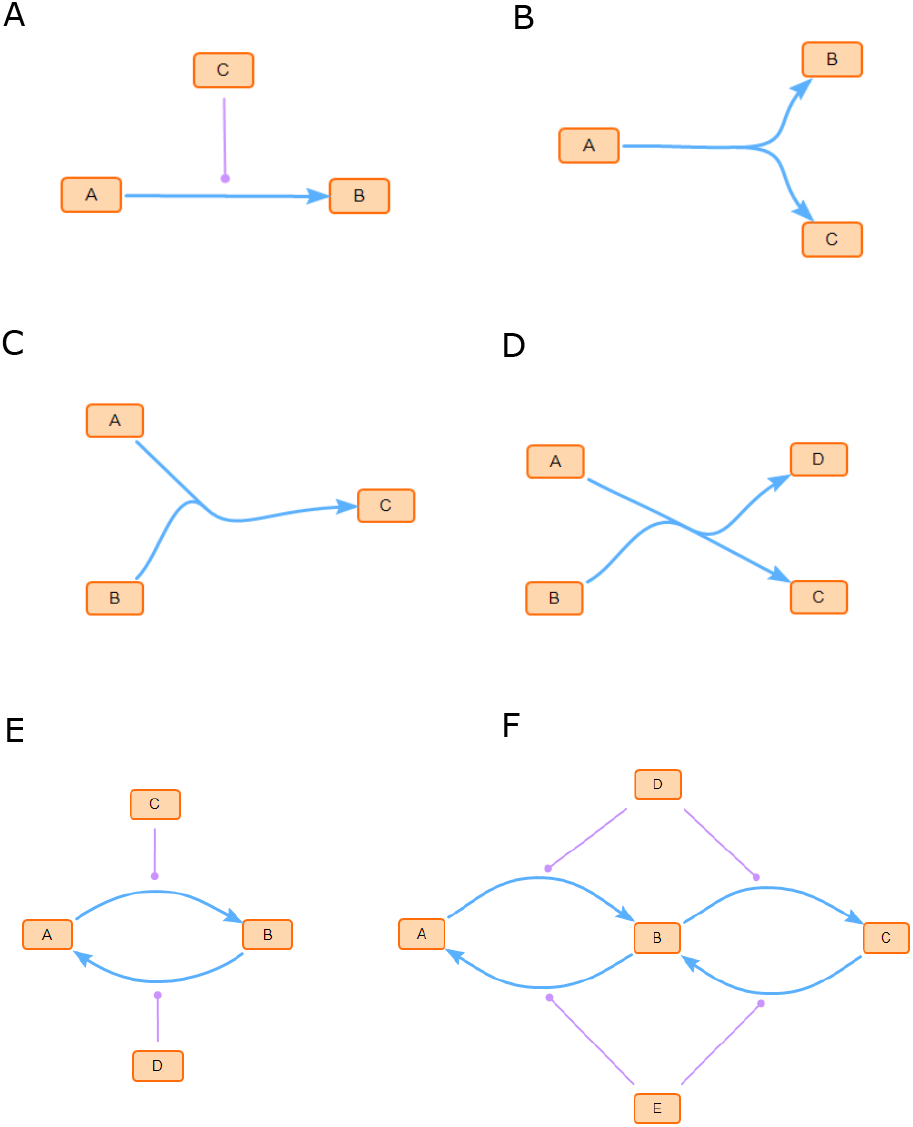
The synthetic random networks were composed of six types of reactions: (A) catalyzed uni-uni, (B, C) binding and unbinding reactions, (D) exchange reactions, (E) catalyzed phosphorylation-dephosphorylation, and (F) dual phosphorylation-dephosphorylation cycles. The figures were generated by SBcoyote version 1.5.0 [28].

Many proteins in signaling networks are found in complexes with other proteins. As a result, the algorithm began by defining a finite set of monomeric proteins and used these in combination with the reaction process shown in Figure 1 to generate a signaling pathway that could consist of multi-protein complexes. Reactions processed from Table 1 were selected at random with predefined probabilities. Each signaling pathway also had a designated input and output species and the network was grown between these two points. The user could also specify the number of species and reactions that should be included in the final network. In this way, arbitrarily complex networks could be generated.

However, not all initially generated networks are viable or useful. Thus, the algorithm imposed structural constraints hat must be followed. These included requiring that all species must be connected to a path in the network that connected the input to the output species, which ensured that no isolated species existed. In addition, this prevented reaction fragments from being isolated from the main body. There were also dynamic constraints in the network generation system. Networks that could not reach a steady state were excluded. We also found that, although some networks appeared complex and connected, perturbations to the input failed to propagate to the output. Because such models could not process information, they were also excluded. We also excluded networks where the input species was directly connected to the output species. We required that there be at least three reactions between the input and output species. The code to generate the synthetic networks was written using the Julia Language (https://julialang.org/) and the simulations were done using the libRoadRunner library [29, 30]; so that we could also export the result as a file with SBML format. See Appendix A for details. There were some configuration settings to generate synthetic signaling networks for biological applications, including the number of species and reactions. See Appendix B for details.

## 3. Results

Figure 2 illustrates two examples of randomly generated signaling networks. Figure 2A included four types of the chemical reaction processes in Table 1. Species were labeled `S1` to `S15`. The reactions included four uni-uni processes: `S6 → S7` catalyzed by `S5, S8 → S6` catalyzed by `S10, S9 → S5` catalyzed by `S15`, and `S11 → S9` catalyzed by `S15`; one bi-bi reactions: `S_in + S14 → S10 + S13`; one single phosphorylation-dephosphorylation cycle: `S12 ⇋ S_out` catalyzed by `S7` and `S1`; and one dual phosphorylation-dephosphorylation cycle `S11 ⇋ S4 ⇋ S3` catalyzed by `S6` and `S2`. Figure 2B included the other two types of the chemical reaction process, namely one uni-bi reaction: `S12 → S12 + S14` and one bi-uni reaction: `S_in + S10 → S11`. As shown, all the random signaling networks had 15 species in addition to input and output species, with 11 reactions. See Figure A.1 for another two samples of synthetic random signaling networks.

**Figure 2:**
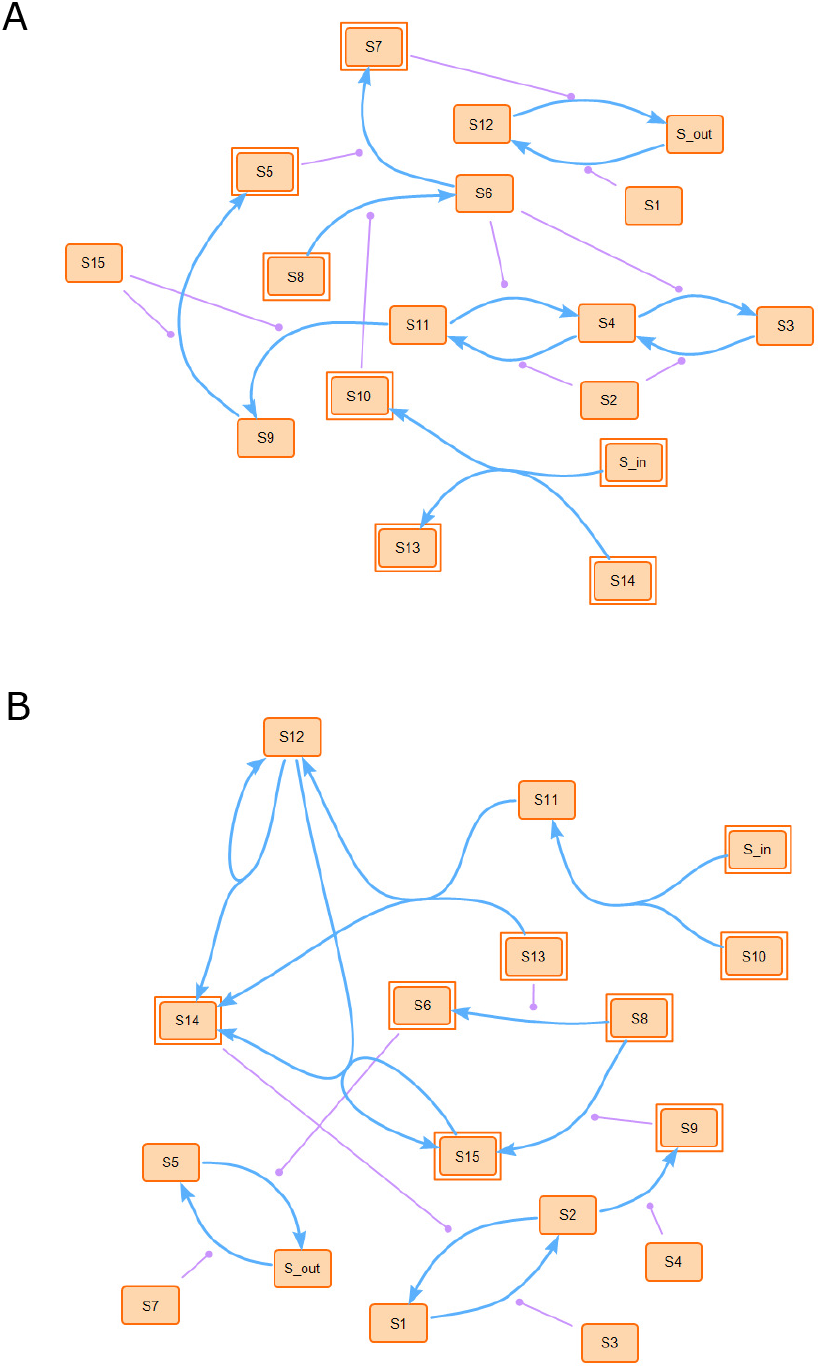
Random signaling networks with 15 species in addition to an input and output species (`S_in` and `S_out`) and 11 catalytic interactions were shown in blue lines with a circle at the end. The figures were generated by SBcoyote version 1.5.0 [28].

The time taken to generate a single network that satisfied the constraints described in the Methods section could range from several minutes to several hours for large networks. The number of trial networks generated to have a single qualified network was from several thousand to several ten thousand depending on the size of the networks. (Table 2).

**Table 2.**
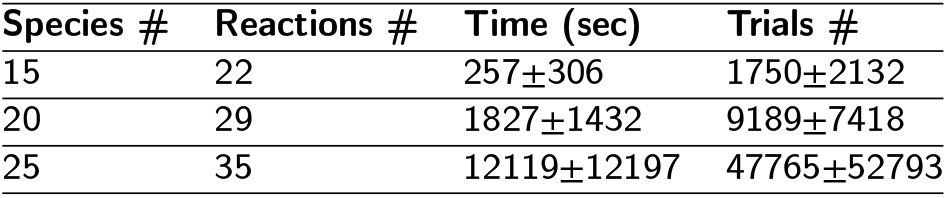
The time taken to generate and find a qualified random signaling network depended on the size of the network. The size of the random network was represented by the number of species involved (input and output species were not included). The errors represented the standard deviations from ten independent runs. All computations reported were done using an Intel i7 9700 processor running at 3.00 GHz with 32GB RAM using Windows 10 with RoadRunner.jl version 0.1.2.

| Species # | Reactions # | Time (sec) | Trials # |
| --- | --- | --- | --- |
| 15 | 22 | 257±306 | 1750±2132 |
| 20 | 29 | 1827±1432 | 9189±7418 |
| 25 | 35 | 12119±12197 | 47765±52793 |

## 4. Discussion

To illustrate the topologies and dynamics of our generated synthetic random signaling networks, we compared them with signaling network models found in the BioModels Database (https://www.ebi.ac.uk/biomodels/). Using the BioModels Database we searched using the category of “Individual Models” and the keywords “signal(l)ing networks”. There were 88 “Manually Curated” signaling models available to download. We downloaded the 88 models and have made them available on GitHub (https://github.com/sys-bio/artificial_random_signaling_network) under `data`. We analyzed these models and found the average species number per model was 27±2 and the average number of reactions per model was 35±3. Therefore, we generated 88 random synthetic signaling networks with 27 species and 35 reactions to compare. The generated 88 synthetic networks are available on GitHub (https://github.com/sys-bio/artificial_random_signaling_network) under `data`.

There were two primary properties of the models: topology and dynamics. From the perspective of topology, we first compared the distribution of reaction degree of species, defined by us as the number of reactions one species belonged to. We also observed the distribution of reaction distance between a random pair of species, defined by us as the minimum number of reactions between the two species. As an extreme case, if two species belonged to one reaction, the reaction distance was zero. Figure 3 showed the two topological distributions of our generated synthetic random signaling networks compared with the signaling networks from the BioModels Database. See Appendix C.1 for code availability. To quantitatively compare the two topological distributions, we calculated the Bhattacharyya distance [31] between the distributions of our generated signaling networks and signaling networks from BioModels. See Appendix C.2 for details. In general, our synthetic signaling networks were comparable with the BioModels signaling networks with Bhattacharyya distances at approximately 0.059 and 0.021 for the reaction degree and reaction distance distributions respectively. See the comparisons with two additional sets of synthetic signaling networks in Figure C.2 and Figure C.3 with small Bhattacharyya distances too. From the perspective of dynamics, we looked at the steady state and oscillatory behavior, shown in Figure 4. Our artificial random signaling networks could generate comparable steady state and oscillatory behavior, which were also commonly observed in the signaling networks from the BioModels Database. Among the 88 signaling networks from the BioModels Database, 56 models showed steady states, and nine models showed dynamics with oscillations. While, among our generated random signaling networks, there was only one case with oscillations.

**Figure 3:**
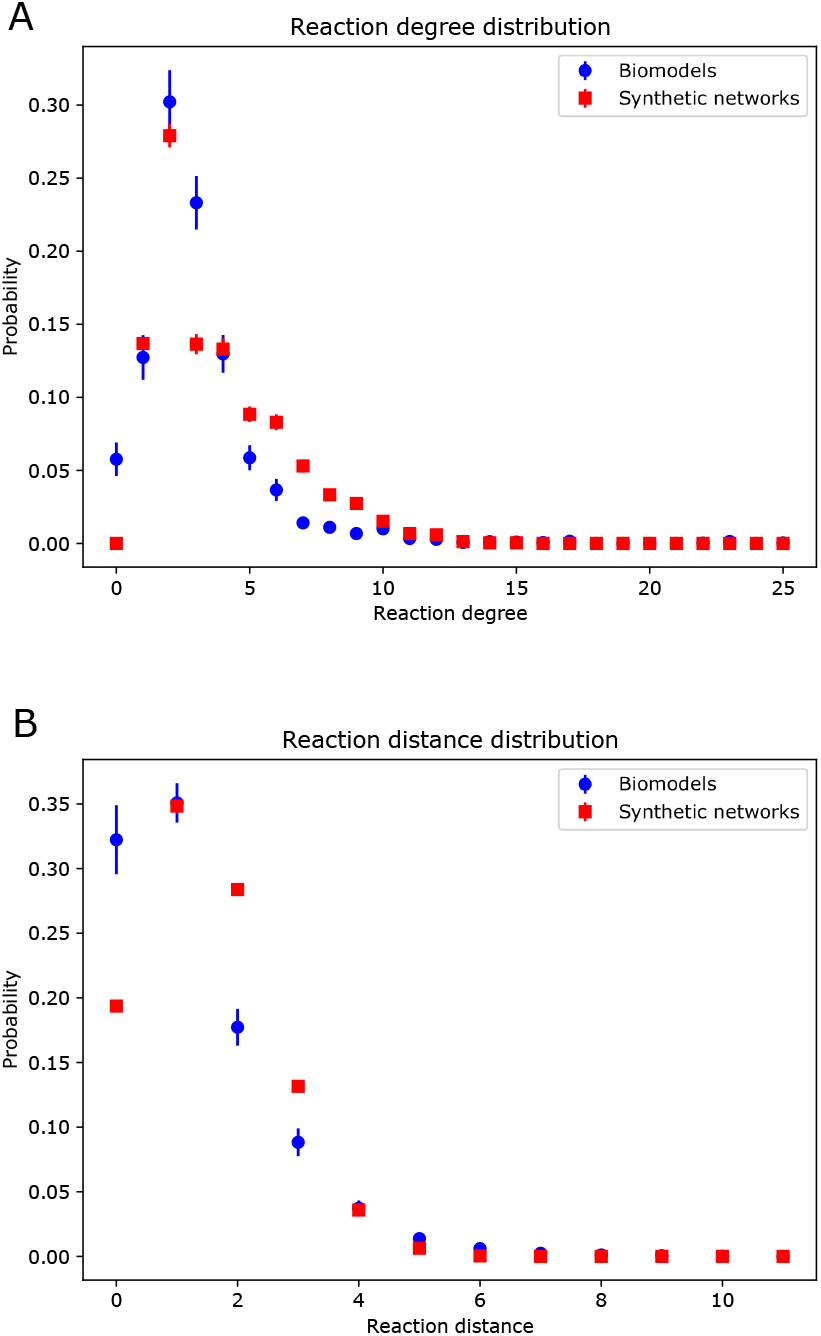
The reaction degree distributions (A) and reaction distance distributions (B) of two sets of signaling network models. The legend “Biomodels” represented the downloaded signaling networks from the BioModels Database. The legend “Synthetic networks” meant the models generated by our method. The error bars represented the standard errors from 88 samples. Our synthetic signaling networks were comparable with the BioModels signaling networks with Bhattacharyya distances at approximately 0.059 and 0.021 for the reaction degree and reaction distance distributions respectively. The distributions were calculated computationally by Python. See code availability in the Appendix C.1.

**Figure 4:**
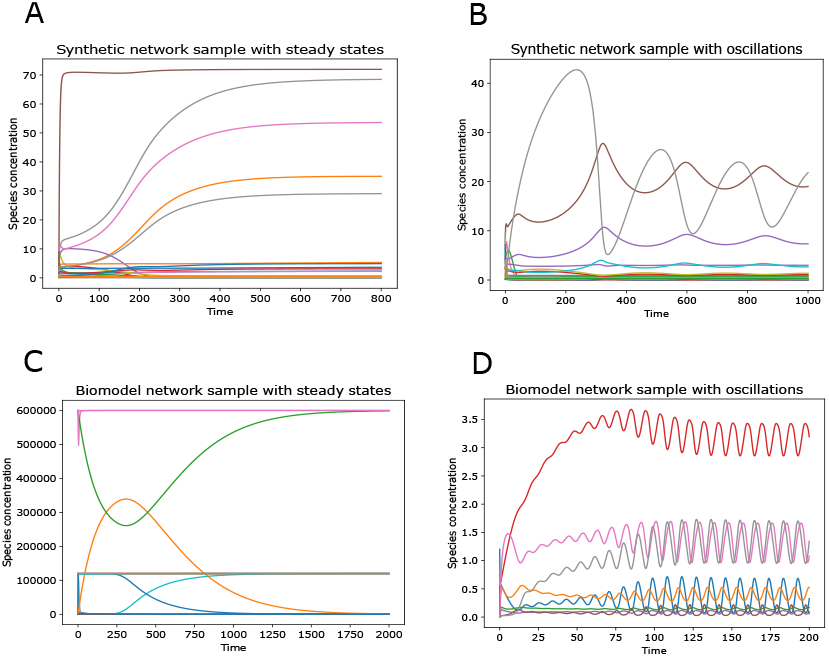
Time-course simulations for four models of signaling networks. The figures showed how the concentrations of species reached their steady state. A and B represented the steady states and oscillations from two sample models of our generated synthetic random signaling networks made up of 27 species (including the input and output species) and 35 reactions. C and D showed the steady states and oscillations from two sample models of signaling networks downloaded from the BioModels Database, i.e., C and D showed the dynamic behaviors of BIOMD0000000427 and BIOMD0000000720 separately. The figures were generated by Tellurium version 2.2.5.2 [21, 22].

There are two existing synthetic random reaction network generating tools, i.e., SMGen [32] and SBbadger [33]. They focus on different perspectives, however, none of the two tools considered the phosphorylation and dephosphorylation cycles which are the critical units in signaling networks. In addition, none of them compared their generated reaction networks with existing databases to discuss the realistic. See the comparison in Table 3.

**Table 3.**
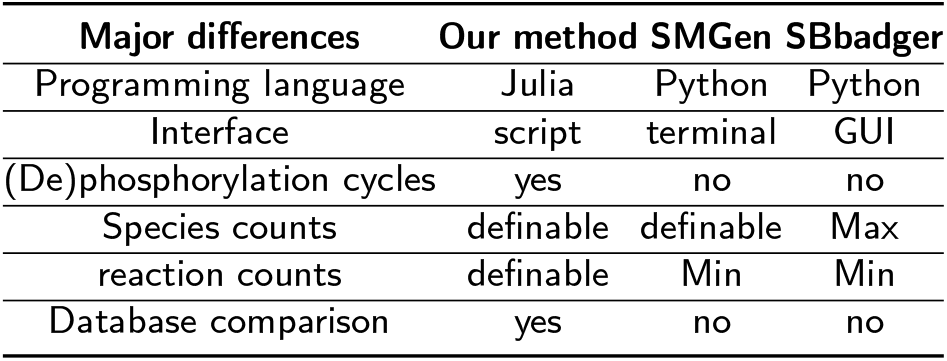
Comparison with two other synthetic reaction network tools.

## 5. Conclusions

The Julia programming language has gained traction with the systems biology community in recent years [34]. Therefore, we developed a Julia script that could allow users to computationally generate artificial random signaling networks to test novel algorithms for parameter fitting, topology mapping, and other analyses. We defined the reaction degree and reaction distance which were applicable definitions from network science to reaction networks, especially to consider enzymes. It would be useful for researchers in related fields to analyze the topology in reaction networks in the future, such as signaling networks or metabolic networks. To illustrate whether our generated signaling networks display meaningful behavior, we compared them with signaling networks from the BioModels Database. The two sets of networks had high similarity from the perspective of topology, namely their reaction degree distributions and reaction distance distributions were similar with low Bhattacharyya distances. In addition, our artificial random signaling networks could generate comparable steady states and oscillations, which were also commonly observed in the signaling networks from the BioModels Database. Therefore, our generated synthetic signaling networks could catch the main features of mechanistic models from the perspectives of topology and dynamics, which could provide sufficient training data or a baseline for validation for future work in the era of AI.

As a comparison with the other two existing tools, SMGen [32] and SBbadger [33], our method considered the critical units of phosphorylation and dephosphorylation cycles which are critical for signaling networks. Additionally, users could define both species and reaction counts within our Julia script. Finally, We compared our generated signaling networks with BioModels Database which was not considered in the other two papers.

## 6. Limitations and future work

The current approach we took has some limitations. The first is that the generated networks did not include any kind of sequestration in the phosphorylation and dephosphorylation cycles. Given the comparable concentrations of substrate and catalytic proteins found in signaling phosphatases, it is now recognized that sequestration [35, 36] can have a significant effect on signaling network behavior [37, 38]. Such effects can’t be modeled with the current version of the software. This would be something to consider for future work.

Another aspect is that we did not examine whether our generated networks had similar graph metrics to networks found in nature. Currently, we only compared our networks to network models found in the BioModels Database. A more thorough investigation would need to be carried out to investigate how the dynamic behavior compares to natural networks. We also did not impose biophysical constraints on the kinetics [39] and parameters. For example, the exact number of the species concentration [40], and the parameters in the kinetic laws [41, 42] have been partially discussed and explored before. Because network behavior is a function of both topology and parameters [43], such constraints could reduce the number of viable topological configurations. Finally, further analysis of which constraints must be either imposed or relaxed to generate realistic network behavior could reveal important principles that real biological networks follow during their evolution.

However, users could apply these synthetic signaling networks for testing methods of network inference [23] to help understanding multi-omics, testing new optimization algorithms [13, 14] and testing how well algorithms can assess uncertainty in fitted parameters [15, 16]. Users could also adjust the values of parameters, i.e., species concentrations and rate constants, according to their future data in hand or demands by themselves following the documentation (sys-bio.github.io/artificial_random_ signaling_network).

## 7. Acknowledgements

This work was financially supported by the National Institute of General Medical Sciences [grant number GM12303201] and the National Cancer Institute [grant number U01CA227544]. The authors are responsible for the content, which does not necessarily represent the opinions of the National Institutes of Health.

We thank Luke Y Zhu for assisting with the development of some of the early code. JX appreciates the valuable comments from the reviewers.

### Appendix A Julia script to generate synthetic signaling networks

The Julia script for generating the synthetic signaling networks is called Ground_truth_generation.jl and is available on GitHub at https://github.com/sys-bio/artificial_random_signaling_network. This Julia script was implemented in Julia 1.6 on Windows 10 with RoadRunner.jl version 0.1.2. See the documentation (sys-bio.github.io/artificial_ random_signaling_network) for using.

**Figure A.1:**
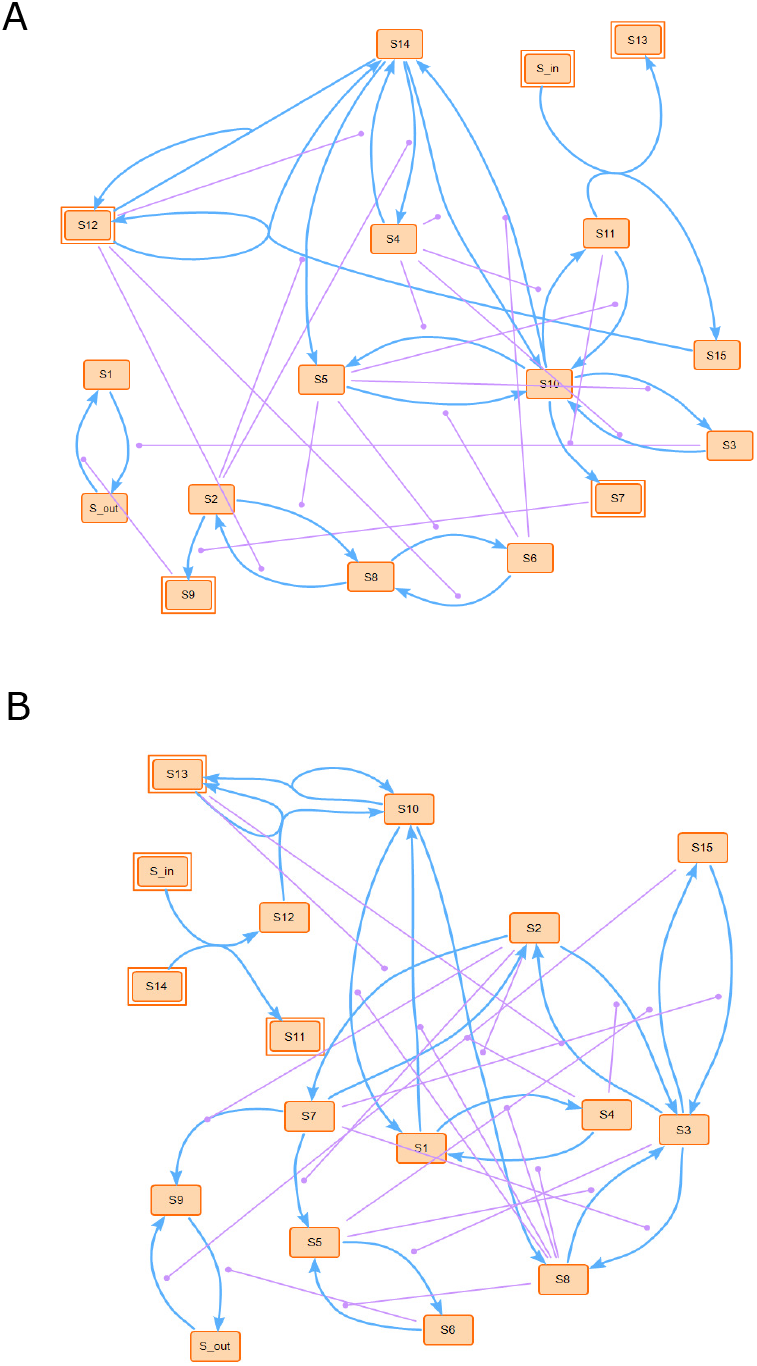
Random signaling networks with 15 species in addition to an input and output species (`S_in` and `S_out`) and 22 catalytic interactions were shown in blue lines with a circle at the end. The figures were generated by SBcoyote version 1.5.0 [28].

Figure A.1 illustrates two more examples of randomly generated signaling networks in addition to Figure 2. Figure A.1(A) included five types of chemical reaction processes in Table 1. Species were labeled `S1` to `S15`. These include three uni-uni processes: `S2 → S9` catalyzed by `S7, S14 → S5` catalyzed by `S2`, and `S10 → S7` catalyzed by `S11`; one bi-uni reaction: `S14 + S12 → S12`; two bi-bi reactions: `S11 + S_in → S15 + S13` and `S12 + S15 → S12 + S14`; two single phosphorylation-dephosphorylation cycles: `S1 ⇋ S_out` catalyzed by `S3` and `S9` and `S4 ⇋ S14` catalyzed by `S2` and `S12`; and three dual phosphorylation-dephosphorylation cycles `S2 ⇋ S8 ⇋ S6` catalyzed by `S5` and `S12, S3 ⇋ S10 ⇋ S11` catalyzed by `S4` and `S5, S5 ⇋ S10 ⇋ s14` catalyzed by `S4` and `S6`. Figure A.1(B) included the 6th type of chemical reaction process, namely one uni-bi reaction: `S10 → S10 + S13`. As shown, all the random signaling networks had 15 species in addition to input and output species, with 22 reactions.

#### Appendix B Adjustable parameters for biological applications

There were some configuration settings in the Julia script file `Ground_truth_generation.jl` that could be adjusted for biological applications including:

- The number of species `nSpeces`, which excludes the input and output species but includes gene species. Therefore, the total number of species should be (`nSpeces`+2).
- The number of gene species `nSpecies_gene`;
- The number of reactions `nRxns`;
- The randomly assigned ranges for species concentrations by `rnd_species_initial` and `rnd_species_range`, for species concentrations in the range of [`rnd_species_initial`, `rnd_species_initial` + `rnd_species_range`);
- The randomly assigned ranges for rate constants by `rnd_parameter_initial` and `rnd_parameter_range`, for rate constants in the range of [`rnd_parameter_initial`, `rnd_parameter_initial` + `rnd_parameter_range`);
- `concentration_perturb` could be used to set the factor that perturbs the concentration at the input species;
- `RXN_MECH_WEIGHT_VALUE` could be used to set the probability of the reaction motifs in the order of uni-uni, uni-bi, bi-uni, bi-bi, single phosphorylation dephosphorylation cycle, and dual phosphorylation dephosphorylation cycle with the sum restriction as 1.
- The number of random networks to generate could be set by changing the variable `sampleSize`.

The Julia script on `Ground_truth_generation.jl` on GitHub at https://github.com/sys-bio/artificial_random_signaling_network is a concrete example with specific biological parameter values shown as below. Users can refer to it to assign their values with their certain demands or based on future biological data in hand.

- `nSpeces` = 15, assigning `nSpeces` as 15, which excludes the input and output species but includes gene species. Therefore, the total number of species should be 17.
- `nRxns` = 22, assigning the number of reactions as 22;
- `nSpecies_gene` = 9, assigning the number of gene species as 9;
- `rnd_species_initial` = 0, and `rnd_species_range` = 10, meaning random generation for species concentrations in the range of [0, 10) [40];
- `rnd_parameter_initial` = 0, and `rnd_parameter_range` = 1, meaning random generation for rate constants in the range of [0, 1) [42];
- `RXN_MECH_WEIGHT_VALUE`set as [0.2, 0.2, 0.2, 0.2, 0.1, 0.1] for the probabilities of reaction motifs in the order of uni-uni, unibi, bi-uni, bi-bi, single phosphorylation dephosphorylation cycle, and dual phosphorylation dephosphorylation cycle with the sum restriction as 1.
- `concentration_perturb` = 2, meaning doubling the concentration at the input species.
- `sampleSize` = 1, meaning generating one sample of synthetic random signaling network.

#### Appendix C Topology analysis

##### C.1 Code availability

The Python scripts for the topology analysis of reaction degree distributions and reaction distance distributions were also available on GitHub at https://github.com/sys-bio/artificial_random_signaling_network under the folder of `topology_analysis`. The code made use of libSBML [44] with a set of SBML files as input and distributions as output. See the section `Topology_analysis` of the documentation (sys-bio.github.io/artificial_random_signaling_network) for using.

##### C.2 Bhattacharyya distance calculations

In statistics, the Bhattacharyya distance is a quantity that represents a notion of similarity between two probability distributions [31]. For probability distributions *P* and *Q* on the same domain *X*, the Bhattacharyya distance is defined as

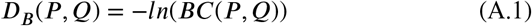

where 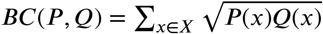 is the Bhattacharyya coefficient for discrete probability distributions. Therefore, the Bhattacharyya distance can measure the distance between two probability distributions, especially applicable to discrete probability distributions.

To judge whether our synthetic signaling networks were comparable with the signaling networks downloaded from the BioModels Database, we measured the Bhattacharyya distances between their distributions of reaction degrees and reaction distances. For all the three comparisons in Figure 3, Figure C.2, and Figure C.3, the results showed that our synthetic signaling networks were comparable with the signaling networks downloaded from BioModels Database with Bhattacharyya distances at approximately 0.06 and 0.02 for the reaction degree and reaction distance distributions respectively.

**Figure C.2:**
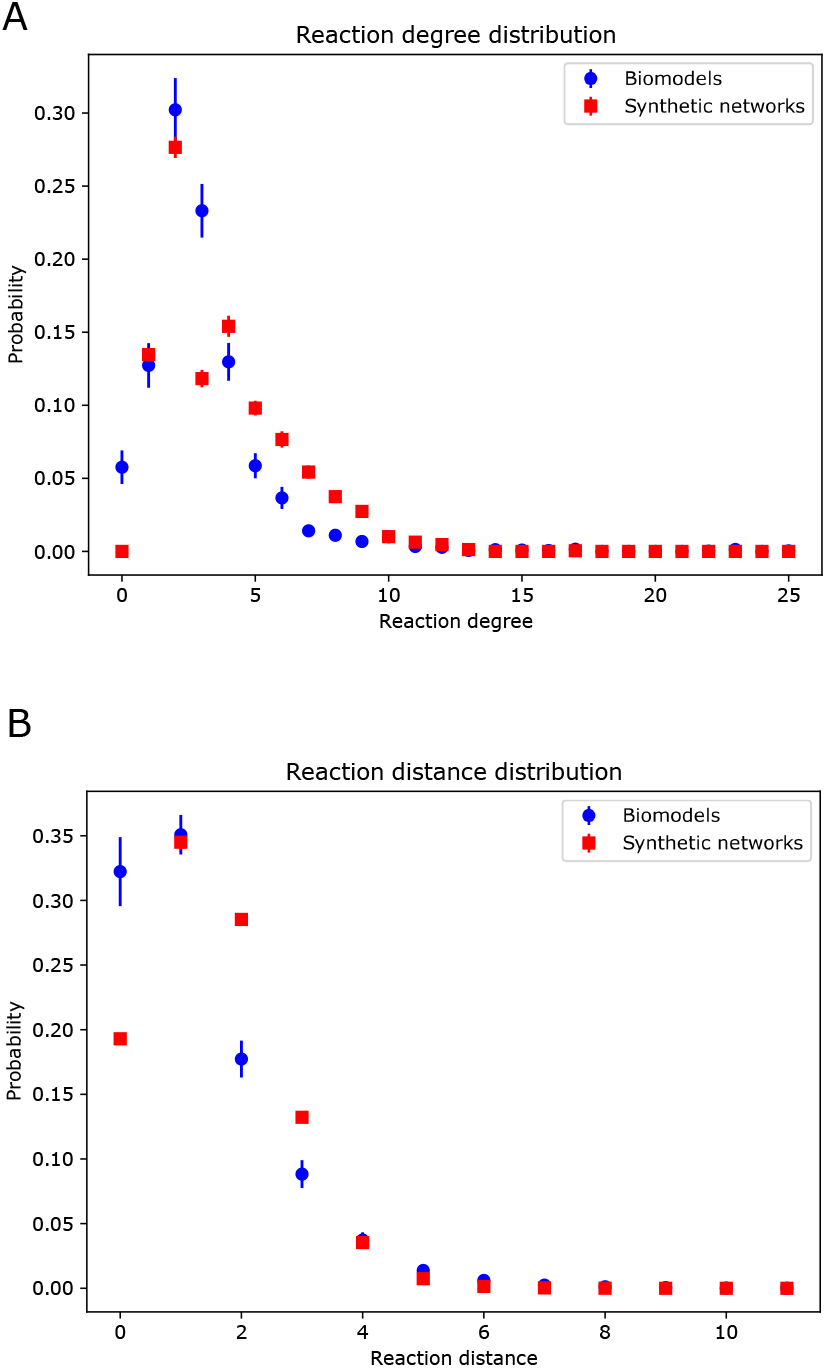
The reaction degree distributions (A) and reaction distance distributions (B) of two sets of signaling network models.

**Figure C.3:**
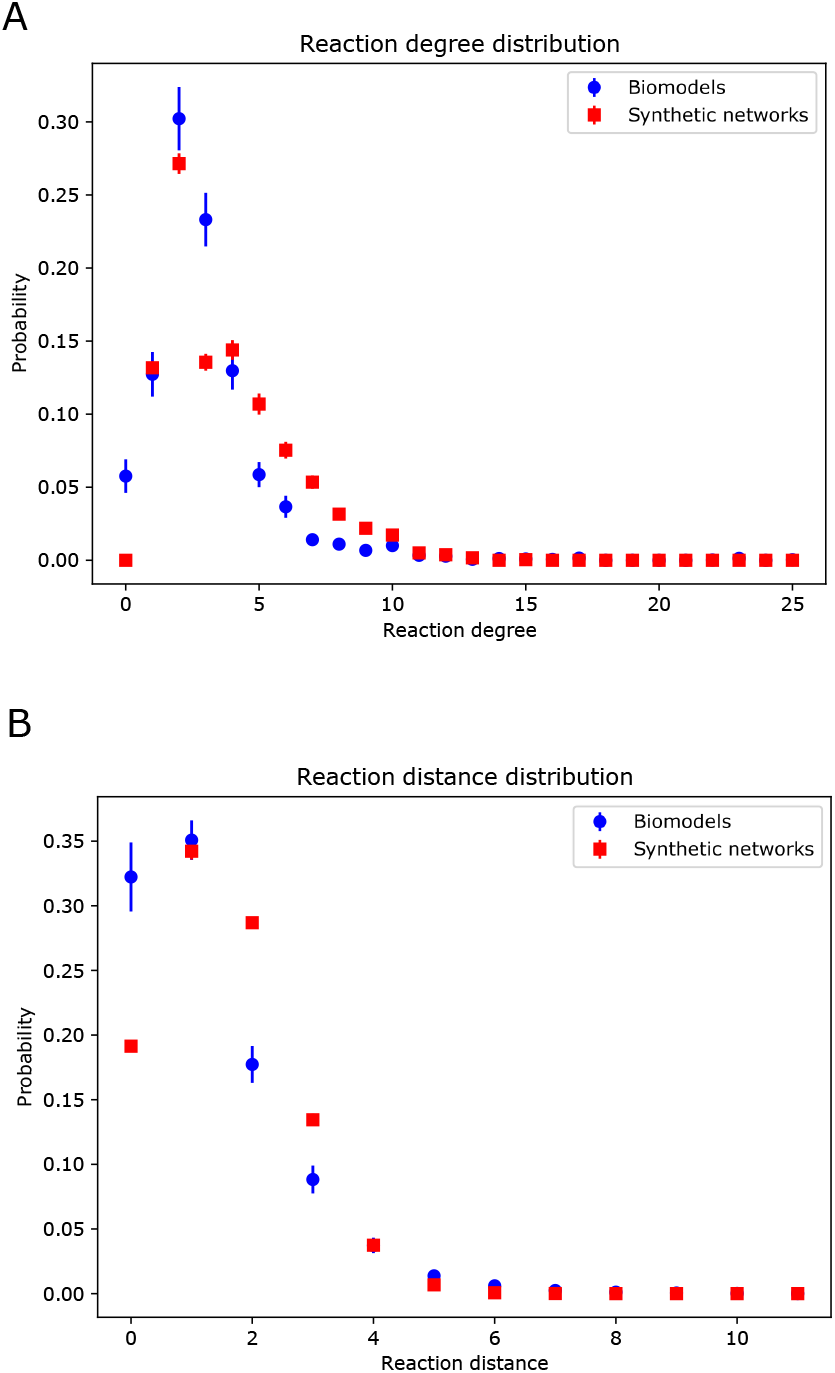
The reaction degree distributions (A) and reaction distance distributions (B) of two sets of signaling network models. The legend “Biomodels” represented the downloaded signaling networks from the BioModels Database. The legend “Synthetic networks” meant the third set of models generated by our method. The error bars represented the standard errors from 88 samples. In general, our synthetic signaling networks were comparable with the BioModels signaling networks with Bhattacharyya distances at approximately 0.059 and 0.021 for the reaction degree and reaction distance distributions respectively.

The legend “Biomodels” represented the downloaded signaling networks from the BioModels Database. The legend “Synthetic networks” meant the second set of models generated by our method. The error bars represented the standard errors from 88 samples. Our synthetic signaling networks were comparable with the BioModels signaling networks with Bhattacharyya distances at approximately 0.064 and 0.020 for the reaction degree and reaction distance distributions respectively.

## CRediT authorship contribution statement

**Jin Xu:** Conceptualization, Data curation, Formal analysis, Investigation, Methodology, Resources, Software, Validation, Visualization, Writing-original draft. **H Steven Wiley:** Funding acquisition, Writing-review and editing. **Herbert M Sauro:** Conceptualization, Funding acquisition, Writing-review and editing.

